# An evolved strain of *Spathaspora passalidarum* produces ethanol from sugarcane bagasse and switchgrass lignocellulosic hydrolysates

**DOI:** 10.1101/2025.09.26.678828

**Authors:** Katharina O. Barros, Kaitlin J. Fisher, Johnathan G. Crandall, Giulia Magni, José Serate, Dan Xie, Yaoping Zhang, Sílvio S. Silva, Trey K. Sato, Chris Todd Hittinger, Carlos A. Rosa

## Abstract

Lignocellulosic hydrolysates, derived from plant biomass, contain various inhibitors that can hinder microbial growth. This study aimed to enable the growth and ethanol production by the xylose-fermenting yeast *Spathaspora passalidarum* in the presence of lignocellulosic hydrolysate inhibitors, particularly acetic acid. Ultraviolet (UV)-induced mutagenesis and adaptive laboratory evolution (ALE) were used to select for mutants with higher tolerance to these inhibitors. The initial mutant strain, MT01, was selected for increased growth in medium containing xylose and acetic acid. This strain underwent further evolution, resulting in the strain ME3.5.5, which showed significant improvements in both growth and ethanol production compared to the parental strain when tested in sugarcane bagasse hemicellulosic hydrolysate (SBHH). Genomic analysis identified non-synonymous and frameshift mutations in four genes, including *CYR1* (encoding adenylate cyclase). These findings suggest that genetically optimized *S. passalidarum* strains could play a crucial role in advancing industrial bioethanol production from lignocellulosic biomass by overcoming the inhibitory effects of compounds found in lignocellulosic hydrolysates.

## Introduction

The production of biofuels from lignocellulosic materials relies on the conversion of carbohydrates, particularly those from cellulosic and hemicellulosic fractions, which mainly include glucose and xylose (Zou et al., 2024). Xylose is the most abundant pentose in hemicellulose, but few microorganisms can ferment this sugar into ethanol. Wild strains of *Saccharomyces cerevisiae*, which are commonly used in industrial alcoholic fermentations, cannot metabolize xylose due to the absence of a xylose assimilation pathway that supports growth (Nalabothu et al., 2023; Ruchala and Sibirny, 2020). The genus *Spathaspora*, which was first described by Nguyen et al. (2006), originally included only a single species, *Sp. passalidarum*, which was isolated from the gut of the beetle *Odontotaenius disjunctus* from the Passalidae family in Louisiana, USA. More recently, this species has been isolated from rotting wood and wood-boring beetles collected in the Brazilian Amazonian rainforest biome (Cadete et al., 2012; Souza et al., 2017) and from rotten wood collected in central China (Ren et al., 2014). Phylogenetic analysis of the complete genome of *Sp. passalidarum* revealed that it belongs to the order Serinales, which decodes CUG codons as serine, instead of leucine. This order also includes the xylose-fermenter *Scheffersomyces stipitis* and opportunistic human pathogen *Candida albicans* (Wohlbach et al., 2011). Currently, *Sp. passalidarum* is one of the most efficient species for converting xylose to ethanol (Cadete and Rosa, 2018), due in part to its possession of genes encoding proteins required for the metabolism of this pentose to ethanol, including a NADH-preferring xylose reductase (Wohlbach et al., 2011).

*Sp. passalidarum* ferments xylose efficiently and achieves high yields of ethanol in rich medium containing xylose as primary carbon source (Cadete and Rosa, 2018; Barros et al., 2024). However, toxic compounds formed during the pretreatment of the lignocellulosic biomass affect the growth and fermentation performance of the species in hydrolysates (Morales et al., 2017).Pretreatment-derived inhibitors include acetic acid, phenolic compounds, and furaldehyde derivatives, which are produced from the hydrolysis of acetyl groups in the hemicellulose fraction, the degradation of lignin, and the dehydration of sugars, respectively. Acetic acid is the major inhibitor present in lignocellulosic hydrolysates (Pattra et al., 2008; Pal et al., 2016; Morales et al., 2017). Its undissociated form goes through the plasma membrane by diffusion and, once within the cell, its proton is dissociated, which leads to the acidification of cytosol (Palmqvist and Hahn-Hägerdal, 2000). To restore the intracellular pH, ATP is consumed to pump protons out of the cell, and the active transport of nutrients is jeopardized, which affects cell growth and fermentation capacity (Guan and Liu, 2020). The combination of acetic acid and other inhibitors creates a synergistic toxicity, in which the harmful effect is greater than the effect caused by only a single compound (Ding et al., 2011; Piotrowski et al., 2014).

The use of lignocellulosic sugars by the xylose-fermenting yeasts for so-called “second generation (2G)” ethanol production depends on strategies that overcome the toxic effects of the inhibitors. To enhance the tolerance of yeasts to inhibitors present in plant hydrolysates, various approaches have been employed, including random mutagenesis, adaptive laboratory evolution (ALE), and genome shuffling (Harner et al., 2015; Morales et al., 2017; Su et al., 2018). These methods are particularly useful because any detoxification of feedstocks for 2G ethanol production will add additional costs (Kumar et al., 2018). Considering that acetic acid is the strongest inhibitor present in sugarcane bagasse hemicellulosic hydrolysate (SBHH) and the synergy by mixing several inhibitors, we hypothesized that ultraviolet (UV)-induced mutagenesis and ALE in medium containing acetic acid as the sole inhibitor would facilitate the adaptation of *Sp. passalidarum* to SBHH, which has several additional toxic compounds, including furfural and 5-hydroxymethylfurfural (5-HMF). Here, we improved the inhibitor tolerance of a Brazilian strain of *Sp. passalidarum*, UFMG-CM-Y469, through UV mutagenesis, followed by ALE in rich medium containing acetic acid and subsequent ALE in SBHH. Finally, we identified de novo mutations by analyzing whole genome sequencing (WGS) data from the ancestral and evolved strains.

## Material and methods

### Yeast strain and UV mutagenesis

*Spathaspora passalidarum* strain UFMG-CM-Y469 (=UFMG-HMD-1.1) was subjected to UV mutagenesis. It was isolated by Cadete et al. (2012) from rotting wood samples collected from Brazilian Amazonian rainforest biome. The strain was obtained from the Coleção de Microrganismos e Células da Universidade Federal de Minas Gerais - UFMG), Belo Horizonte, Brazil.

The yeast strain was streaked on yeast malt agar (1% glucose, 0.5% peptone, 0.3% yeast extract, 0.3% malt extract, and 2% agar). After 48 h, a single colony was propagated into 15-mL tubes containing 5 mL of YPX (2% xylose, 2% peptone, and 1% yeast extract). The tubes were incubated in an orbital shaker under 200 rpm, at 30ºC for 24 h. The cells were recovered and washed with sterile deionized water, and the cell concentration was adjusted to 1 x 10^7^ cell/mL for each strain. To determine the survival curve, the cultures were subjected to a series of 10-fold dilutions. One hundred microliters of each dilution were inoculated in YPX agar plates. The plates were placed in a Crosslinker CL-1000 (UVP, England), which produces UV radiation at 365 nm, and they were exposed to several intensities of UV radiation (1,000; 1,500; 2,000; 3,000; 4,000; and 5,000 μJ/cm^2^). The plates were incubated at 30ºC for 48 h, and the survival rate was calculated as previously described (Hou and Yao, 2012). After identifying the optimal UV intensity using the survival curve, which reflects the level of UV radiation that reduces cell viability to a target percentage (typically around 5 to 20%) (Lawrence, 2002), mutagenesis was carried out according to Hou and Yao (2012). This range ensures that, although a significant portion of cells are killed, enough survive to allow for the accumulation of random mutations.

Cells were pre-cultured overnight in YPX, and the cell concentration was adjusted to 1 x 10^7^ cell mL^-1^. One hundred microliters were spotted onto YPXAC (2% xylose, 2% peptone, 1% yeast extract, and 0.25% acetic acid) agar plates. Plates were subjected to 1,500 μJ/cm^2^ UV radition, covered with aluminum foil immediately after the exposition of UV to avoid photoreactivation repair, and incubated at 30ºC. After 96 h, the colonies were transferred to YPXAC plates by replica plating and incubated at 30ºC for 48 h. Then, the colonies were transferred to YPX agar plates and, after 48 h, plated back to YPXAC. Single colonies were grown in GYMP broth (2% glucose, 0.5% yeast extract, 0.5% malt extract, and 0.2% Na_2_PO_4_) and stored in 20% glycerol at −80°C. Three independent UV mutagenesis assays were conducted.

### Microtiter-plates growth curves and shake flasks fermentation assays in YPXAC and SBHH

To compare the growth in acetic acid of the parent and mutants, the strains were cultured overnight into 5 mL of YPX. After pre-culture, cells were recovered and washed with sterile water, and an aliquot of each strain was transferred to a 96-well plate, which contained YPX medium with and without 2 g L^-1^ of acetic acid, at the initial OD_600_ of 0.1. The final volume was 250 μL. Medium without inoculation was used as negative control. The plate was placed in Tecan (Infinite, Switzerland) at 30ºC. Absorbance at 600 nm was monitored every 30 min for 144 h. Kinetic parameters were calculated with R package growth rates in R v3.6.3 using the RStudio v1.3.1073 platform.

Mutants selected based on growth curves, along with the parent strain, were evaluated in fermentation assays with varying concentrations of acetic acid (2 and 3 g L^-1^). The strains were pre-cultured in YPX for 16 hours, and approximately 0.5 g/L of cells were inoculated into 125 mL shake flasks containing 50 mL of YPXAC (with either 0.2% or 0.3% acetic acid and 3% or 5% xylose). Samples were taken at regular intervals every 2 hours for the first 12 hours in the medium containing 2% acetic acid. For the medium containing 3% acetic acid, samples were taken every 24 hours for 72 hours. The samples were then centrifuged at 13,500 x g for 10 minutes. Supernatants were stored at −20ºC. Cell growth was determined by correlating optical density or OD_600_ (Biospectro SP-22, São Paulo, Brazil) with the cell dry weight (CDW). Concentrations of xylose, ethanol, and xylitol were determined by high-performance liquid chromatography or HPLC (Shimadzu, Kyoto, Japan) using the following conditions: Supelco Analytical C-610 H column (Sigma-Aldrich, USA), maintained at 45°C; volume injection of 20μL; refractive index detector RID 10-A (Shimadzu, Kyoto, Japan); and 5 mM H_2_SO_4_ mobile phase as eluent at a flow rate of 0.6 mL min^-1^. Graphs were constructed using the ggplot2 package in R v3.6.3 using the RStudio v1.3.1073 platform.

### Lignocellulosic hydrolysates

Sugarcane bagasse hemicellulosic hydrolysate (SBHH) was obtained by acid hydrolysis in a 250-L bioreactor at 120°C for 20 min in 98% sulfuric acid at a ratio of 1:10 (i.e. 100mg H_2_SO_4_ per gram of sugarcane bagasse). The liquid fraction was recovered by filtration, the pH was adjusted to 5.5 with calcium oxide (CaO), and it was autoclaved at 111ºC for 15 min. The final concentrations in SBHH of sugars and inhibitors were determined by HPLC (Barros et al., 2022). Detoxification and supplementation were not performed.

Switchgrass hydrolysate (ASGH) was produced by Ammonia Fiber Expansion (AFEX) pretreatment and enzyme-loading method (Serate et al., 2015). Biomass was loaded into tubes for hydrolysis to reach the final content of 7% glucan to obtain a hydrolysate with ∼60 g L^-1^ glucose.Water was added to the 7% glucan loading AFEX-pretreated switchgrass biomass, and it was autoclaved for 2 h at 121ºC. Then, the pH was adjusted with HCl (∼37–38% HCl, w/v) and a mixture of cellulase (NS 22257) and xylanase (NS 22244) from Novozymes (Franklinton, NC, USA) was added. The total enzyme loadings were 80 mg protein g^-1^ glucan of biomass for cellulase and 13 mg protein g^-1^ glucan of biomass for xylanase. After 2-4 days of mixing, the hydrolysis was carried out at 50 °C with stirring at 700 rpm for 7 days. Solids were removed by centrifugation at 8200 × *g* at 4 °C for 10–12 h, the supernatant was filtered through 0.5 μm GVS Maine Glass Prefilters (Thermo Fisher Scientific Inc. Waltham, MA, USA) and 0.2 μm Filter Units (VWR International, Radnor, PA, USA), and the ASGH was stored at 4°C (Zhang et al., 2020).

### Adaptive laboratory evolution (ALE) experiments

The selected mutant MT01 underwent ALE. The strain was evolved aerobically in 50 mL shake flasks containing 20 mL of YPXAC medium (2% xylose and 0.15% acetic acid) at 30°C with shaking at 200 rpm. Every 24 hours, a 100 μL aliquot was transferred to fresh medium. After five passages at a given acetic acid concentration, the concentration was increased by 0.5 g L^-1^, continuing this process until reaching 3.5 g L^-1^ of acetic acid. During the ALE experiment, cultures from each passage were diluted to check the viability in YPXAC agar plates, and single colonies were isolated and cryopreserved. The ALE experiment in YPXAC was followed by ALE using SBHH. This ALE experiment started with a culture of ME3.5.5 from the last passage in YPXAC. The evolved strain was inoculated in 50mL-shake flasks containing 20 mL of medium (70% sterile water and 30% SBHH), and the dilution was decreased by 10% in each round. This process was repeated until the SBHH concentration was 60% (v/v), with direct transfer to a new concentration of hydrolysate.

### Fermentation assays in YPX, SBHH, and ASGH

Evolved strains were inoculated in YPX medium, and the flasks were incubated at 30°C in an orbital shaker at 200 rpm for 16 h. The cells were recovered as described above and suspended in YPX (5% xylose, 2% peptone, and 1% yeast extract), SBHH, or ASGH. The fermentation was performed in 125 mL shake flasks with 50 mL of medium and incubated at 30°C and 200 rpm. The fermentation was monitored by collecting samples every 24 h for fermentations in YPX and SBHH and every 6 h for ASGH. The samples were stored at −20°C until analysis. The cell concentrations were determined by measuring the OD_600_. After determining the cell concentration, the samples were centrifuged at 2600 × *g* for 10 min, and the supernatant was stored at −20°C for analysis. The experiment was performed in triplicate.

Xylose, glucose, xylitol, ethanol, and acetic acid levels were determined using HPLC (Shimadzu, Kyoto, Japan) with a Supelcogel C610H ion exclusion column (Sigma-Aldrich, USA) at 45°C and a refractive index detector RID-10A (Shimadzu, Kyoto, Japan). The mobile phase was 5 mM H_2_SO_4_ at a flow rate of 0.6 mL min^-1^ (Barros et al., 2024).

### Genome sequencing

To confirm that all the mutant and evolved strains were still *Sp. passalidarum*, D1/D2 sequences of large subunit (26S) of ribosomal DNA of all selected strains were performed (Kurtzman and Robnett, 1998). Later, we extracted genomic DNA from the strains UFMG-CM-Y469, ME3.5.5, and MEH30.1 using standard phenol:chloroform extraction and Illumina library preparation methods (Kominek et al., 2019). Briefly, cells were grown to saturation in YPD (1% yeast extract, 2% peptone, and 2% glucose) broth, collected by centrifugation with approximately 500 mL 0.5 mm acid-washed beads (Sigma #G8772), and resuspended in DNA lysis buffer (10 mM Tris, 1 mM EDTA, 100 mM NaCl, 1% SDS, 2% Triton X-100 in water). Samples were extracted twice with 25:24:1 phenol:chloroform:isoamyl alcohol (Sigma #P2069), precipitated overnight at −80ºC in 100% ethanol, collected by centrifugation, washed twice with 70% ethanol, dissolved in 10 mM Tris-Cl (pH 8), and treated with RNase A (VWR #97064-064) for 30 min at 37ºC. Libraries were prepared with NEBNext Ultra DNA Library Prep kit Illumina (NEB #E7370L), which was performed according to the manufacturer’s protocol. Libraries were submitted for 2×150 bp sequencing on an Illumina NovaSeq 6000 instrument.

### Analysis of variants

Adapters were first trimmed from de-multiplexed paired end reads using Trimmomatic (Bolger et al., 2014). Trimmed reads were then aligned to a *Sp. passalidarum* reference assembly (accession #AEIK00000000, Wohlbach et al. 2011) using BWA-MEM (Li, 2013). Variants were called using FreeBayes v1.3.1 (Garrison and Marth, 2012) using standard filters. Resulting VCF files were filtered for shared and unique variants using VCFtools. Candidate mutations were manually confirmed in IGV. Genes containing mutations were identified first by BLASTing the mutated region to the annotated reference genome (AEIK00000000). Nucleotide annotations matching the mutated region were then BLASTed against annotated *S. cerevisiae* proteins using BLASTX. *S. cerevisiae* proteins with significant (e-value ≤ 1e^-5^) scores considered homologous.

Genomes were independently examined for structural variants using samtools depth (Li and Durbin, 2009). Euploidy of all genomes was confirmed by dividing median chromosome coverage by median genome-wide coverage for each chromosome. Structural variants unique to evolved strains were screened through visual inspection of chromosome coverage plots.

## RESULTS

### *Sp. passalidarum* mutants

We selected a UV intensity of 1500 µJ/cm^2^, which resulted in the survival of 9% of cells of *Sp. passalidarum* UFMG-CM-Y469. After UV radiation, cells were transferred to new YPX plates and then transferred to YPXAC by replica plating (Fig. 1a). This procedure was repeated three times. At the end, 55 colonies were recovered from YPX plus 0.25% acetic acid plates. The total colonies were obtained from the three independent UV mutagenesis experiments. The size of the colonies varied from large to petite. After the last transfer, petite colonies were reduced significantly.

**Figure 1.**
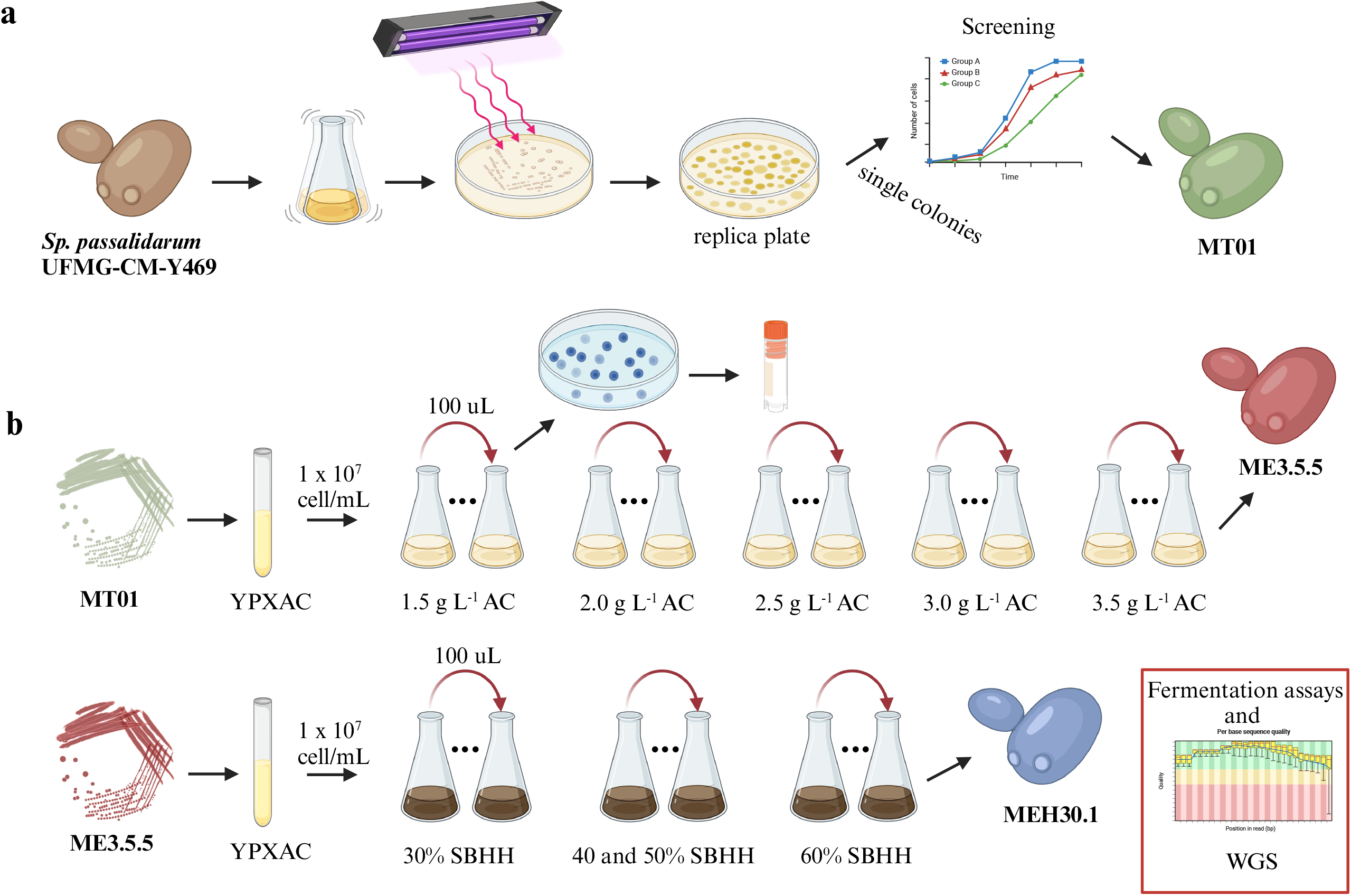
Diagram of a) UV mutagenesis experiment and b) adaptive evolution of MT01. Abbreviations: WGS - whole genome sequencing, YPXAC - yeast extract, peptone, xylose, and acetic acid. Figure was generated in Biorender (https://app.biorender.com/).

Six mutants (MT01, MT02, MT37, MT53, MT87, and MT101) showed enhanced cell growth compared to the parental strain. A triplicate shake flask experiment was performed to compare those mutants with the parental strain in YPX and YPXAC to verify whether the mutants were able to produce ethanol and still ferment xylose in both conditions. MT01 was the only mutant to exhibit growth, xylose consumption, and ethanol production superior to the parental strain in YPXAC (Fig. 2). From 30 g L^-1^ of xylose, MT01 consumed 41% of the sugar in 12 h, and it produced 5.31 g L^-1^ (0.43 g g^-1^) of ethanol and 2.30 g L^-1^ of biomass (Fig. 2a). This mutant produced 43% more ethanol than parental strain UFMG-CM-Y469, which reached 3 g L^-1^ of ethanol and 2.13 g_CDW_ L^-1^ of biomass while consuming 34% of xylose in YPX plus 2 g L^-1^ of acetic acid. MT01 achieved a maximum ethanol yield of 0.46 g/g (13 g/L), which is close to the theoretical maximum yield of 0.51 g/g. Additionally, the strain consumed 95% of the sugar within 10 to 12 hours.

**Figure 2.**
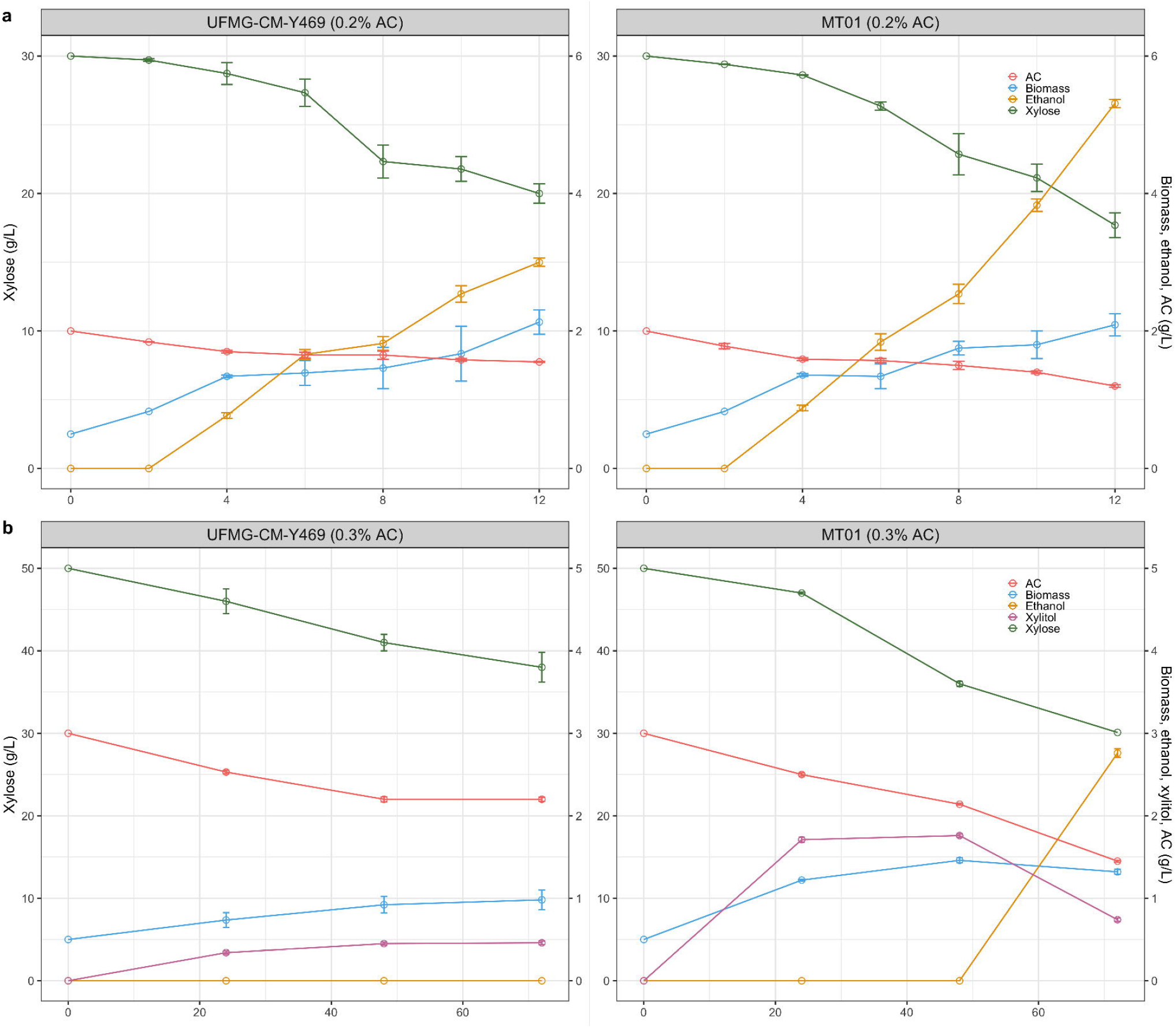
Shake flask fermentations of (a) YPXAC (2 g L^-1^ acetic acid (AC)) in 12 h and (b) YPXAC (3 g L^-1^ acetic acid) in 72 h by *Sp. passalidarum* MT01 (right) and parental strain UFMG-CM-Y469 (left). Error bars indicate the standard deviation from triplicate shake flasks.

In agar plates containing acetic acid, the growth of UFMG-CM-Y469 was limited to 1.5 g L^-1^ of acetic acid. This growth was extended in the liquid medium plus inhibitor, and it still fermented xylose and produced ethanol in 2 g L^-1^ of acetic acid. Increasing this concentration to 3 g L^-1^, the parental strain consumed 12 g L^-1^ of xylose, but ethanol production was negligible, and little xylitol was excreted (Fig. 2b). The mutant MT01 produced 2.76 g L^-1^ of ethanol in 72 h, but it also excreted xylitol. Both strains consumed acetic acid.

Mutants MT02, MT37, MT53, MT87, and MT101 did not show desirable phenotypes (i.e. enhanced ethanol production in shake flask experiments). MT37 and MT101 lost the ability to ferment xylose, while MT02, MT53, and MT87 had a similar or inferior performance compared to the parental strain UFMG-CM-Y469. Since the parental strain was not able to grow in SBHH and produce any fermentation products, we only tested MT01 in SBHH to check whether the mutant would grow, but no growth was observed.

### A further evolved strain was able to grow and produce ethanol in SBHH

Since the mutant MT01 showed enhanced ethanol production in the presence of acetic acid compared to the parent strain, it was chosen for an additional round of ALE. We adapted MT01 over many generations (∼ 300) until the concentration of 3.5 g L^-1^ acetic acid (Fig. 1b) was reached, yielding the strain ME3.5.5, which showed ethanol production and cell growth in the hydrolysate. The strain ME3.5.5 was then subjected to adaptive laboratory evolution (ALE) using SBHH hydrolysate, with the hydrolysate concentration gradually increased from 30% to 60%. The hydrolysate used in this work was prepared with the hemicellulosic fraction from the sugarcane bagasse. Its composition was 2% xylose, 0.2% glucose, 0.45% acetic acid, 0.013% HMF, and 0.02% furfural. The ALE experiment was carried out with a maximum of 60% of the hydrolysate, but we also tested the evolved strains without diluting the hydrolysate. The evolved strain was able to grow and produce fermentation products in SBHH. In the triplicate shake flask experiments, ME3.5.5 obtained from ALE in YPXAC reached its maximum ethanol yield after 72 h of fermentation at 0.42 g g^-1^ (9 g L^-1^) (Fig. 3a). The complete utilization of xylose and acetic acid by the evolved strain occurred in 72 h and 96 h of fermentation, respectively. As the glucose was present at such low amounts in this hydrolysate, glucose was rapidly depleted, and the yeasts were able to start consuming xylose in the first 24 h. ME3.5.5 presented a maximum specific growth rate (μ_max_) of 0.05. It was also tested in YPX to assess potential trade-offs (see supplementary Fig. S1). Starting with 50 g/L of xylose, the yeast produced high amounts of ethanol, and nearly all the xylose was consumed within 24 hours with a μ_max_ of 0.12 h^-1^. The resulting strain from ALE with SBHH, MEH30.1, exhibited growth identical to that of ME3.5.5. As there were no phenotypic improvements (e.g. etanol production and/or xylose consumption) observed in MEH30.1 in SBHH 5 (see supplementary Fig. S2), we did not conduct further analyses with this strain.

**Figure 3.**
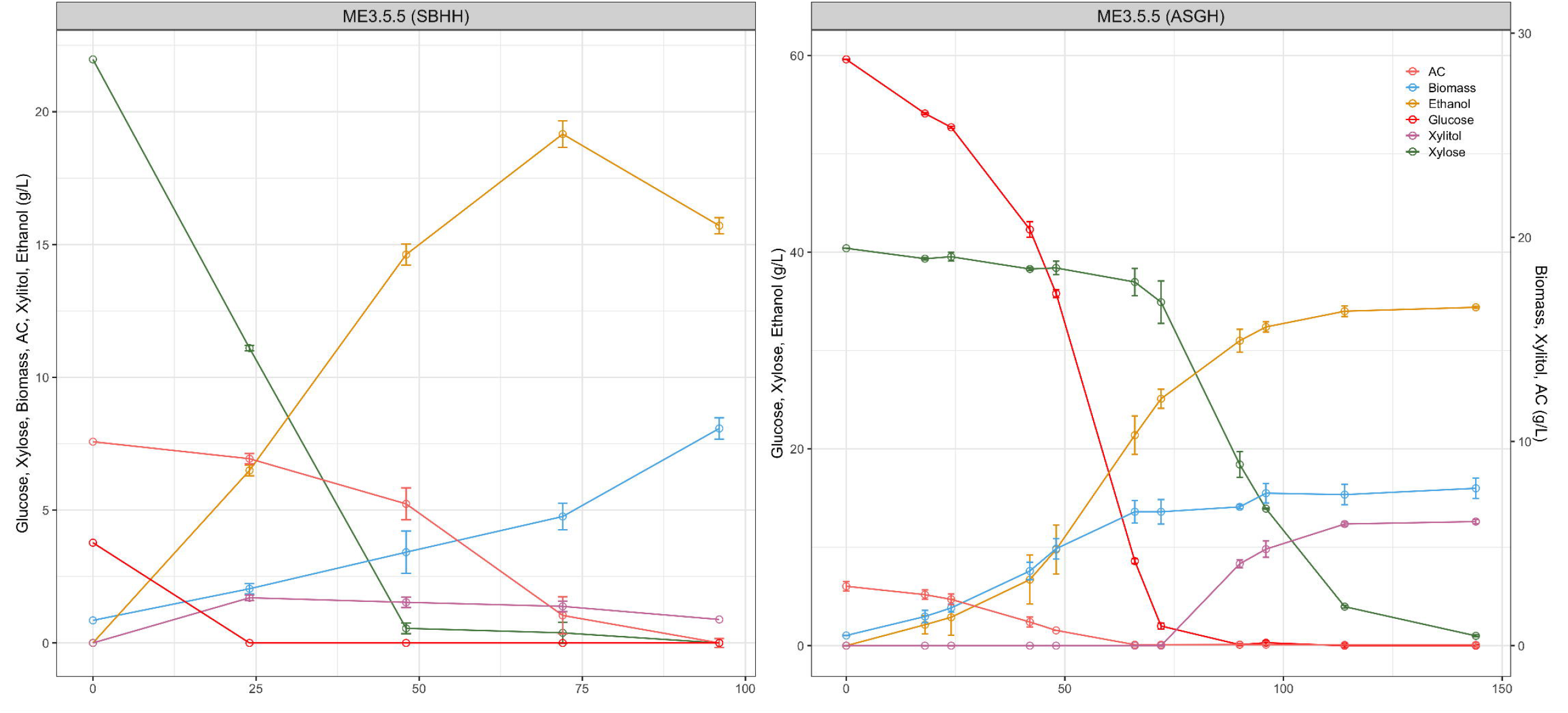
Shake flask fermentations of SBHH and ASGH by evolved strain ME3.5.5 from ALE with YPXAC. Error bars indicate the standard deviation from triplicate shake flasks.

Since we used hemicellulosic hydrolysate, other sugars (e.g. arabinose, mannose, cellobiose, and galactose) in addition to xylose and glucose were also likely present. Unfortunately, it was not possible to measure the other sugars present in the SBHH. This analytical limitation is a possible explanation for the continued production of ethanol after xylose and glucose depletion.

### Performance of evolved strains in ASGH

Since the SBHH is produced from hemicellulosic fractions relevant to Brazilian industry, we also decided to test the native and evolved strains in a switchgrass hydrolysate prepared with both hemicellulosic and cellulosic fractions of a lignocellulosic material. ASGH is produced by Ammonia Fiber Expansion (AFEX) pretreatment of switchgrass, followed by enzymatic hydrolysis using cellulase and xylanase from Novozymes (Franklinton, NC, USA) to release fermentable sugars. ASGH has a higher content of glucose and xylose, about 60 g L^-1^ and 40 g L^-1^, respectively, than SBHH. On the other hand, some inhibitors were present in lower amounts: 2.8 g L^-1^ acetic acid and 0.04 g L^-1^ furfural. The evolved strain had similar growth of the parental strain UFMG-CM-Y469 in this hydrolysate (Fig. 4), and it showed similar values for fermentation products, biomass, and consumption of sugars and acetic acid. Negligible amounts of xylose were used during the first 72 h of fermentation. As expected, the pentose was fully consumed after 72 h, when glucose and acetic acid were also depleted. The maximum ethanol yield was 33.6 g L^-1^ after 144 h. But the maximum productivity, 0.329 g L^-1^ h^-1^, took place at 90 h of fermentation with an ethanol yield of 0.498 g g^-1^ (29.6 g L^-1^). At this time, the yeasts started to accumulate xylitol. The strains reached about 8 g L^-1^ of biomass with μ_max_ of 0.14 h^-1^.

### Mutations identified by whole genome sequencing

To evaluate candidate genetic bases of the adaptation, we performed whole genome sequencing on UFMG-CM-Y469 and the evolved strains ME3.5.5 and MEH30.1. Depth plots revealed no aneuploidies or large CNVs in the parental or evolved strain. Using a variant-calling pipeline, we identified two types of mutations: single-nucleotide polymorphism (SNPs) and small insertions and deletions (indels). First, we looked for mutations shared by the two evolved strains, which were absent in the UFMG-CM-Y469 ancestor. Five heterozygous mutations were identified among the strains, one intergenic and four non-synonymous mutations (Table 1). A C583T mutation resulting in a nonsynonymous L195F change was found in a gene encoding a poorly conserved hypothetical protein (XM_007377149). The remaining three protein-coding sequence mutations are all predicted to disrupt the coding sequences of genes with homologs conserved in *S. cerevisiae*. A C1208T missense mutation, resulting in a P403L non-synonymous substitution, was found in the *Sp. passalidarum* homolog of *ERT1* (XM_007373880.1); a C492A mutation causing a Y143* nonsense mutation, resulting in a premature stop codon, was found in the homolog of *NPL4* (XM_007372648); and two base pair deletion between nucleotides 3009 and 3011 in the *CYR1* causing a D1003 frameshift. Since we did not sequence MT01, we could not differentiate between UV-derived mutations underlying improved performance in MT01 and spontaneous mutations acquired by ME3.5.5.

**Table 1:**
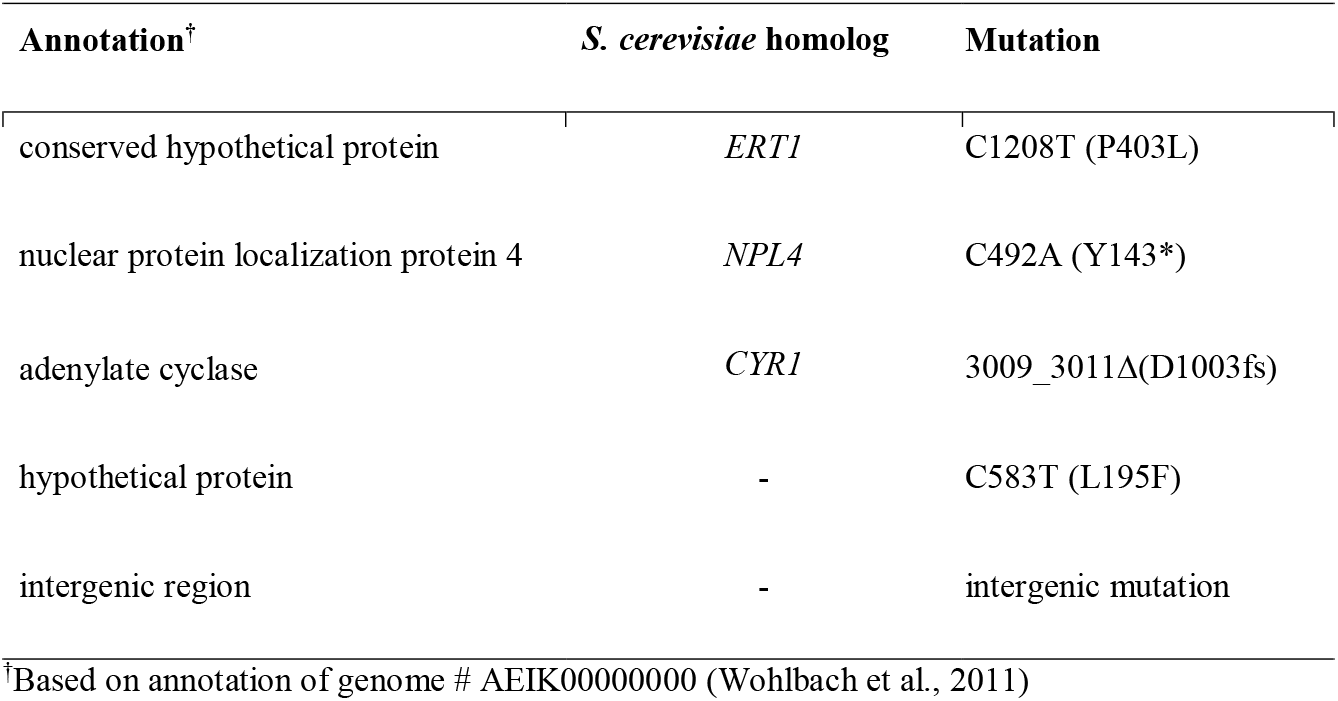
Five heterozygous mutations shared by the evolved strains and absent in the ancestor.

| Annotation<sup>†</sup> | *S. cerevisiae* homolog | Mutation |
| --- | --- | --- |
| conserved hypothetical protein | *ERT1* | C1208T (P403L) |
| nuclear protein localization protein 4 | *NPL4* | C492A (Y143*) |
| adenylate cyclase | *CYR1* | 3009_3011Δ(D1003fs) |
| hypothetical protein | - | C583T (L195F) |
| intergenic region | - | intergenic mutation |

<sup>†</sup>Based on annotation of genome # AEIK000000000 (Wohlbach et al., 2011)

## Discussion

A major challenge in biofuel production from lignocellulosic hydrolysates is the generation of toxic compounds during pretreatment and hydrolysis, which can inhibit microbial metabolism. Notably, the synergistic effects of acetic acid with furfural and phenolic compounds inhibit amino acid biosynthesis and carbon catabolism, leading to significant decreases in pyruvate and succinate levels (Ding et al., 2011). Comparisons of these inhibitors individually have shown that acetic acid has the most detrimental effects on yeast growth and metabolism (Morales et al., 2017; Ding et al., 2011). Developing microorganisms with enhanced tolerance to acetic acid and other toxic compounds could advance our understanding of traits of biotechnological interest, which could then be genetically manipulated for improvement.

In this study, a strain of *Sp. passalidarum* was subjected to UV mutagenesis, followed by adaptive laboratory evolution (ALE). The first phenotypic trait observed after UV exposure was an increase in colony size. Larger colonies suggest a primary resistance mechanism to mitigate UV damage, as their increased size allows them to maintain greater reserves of water and other resources (Goldman and Travisano, 2011). Additionally, larger colonies may be associated with increased lipid metabolism efficiency, possibly as a compensatory mechanism for enhanced lipid synthesis (Wang et al., 2009). Conversely, petite colonies, which are often linked to respiratory deficiency (RD) mutations, were more prevalent on YPXAC plates following mutagenesis, but these were not selected for further study. This prevalence likely results from UV-induced mutations in mitochondrial DNA, which impair respiratory function (Stewart, 2017).

A mutant MT01 was selected to be subjected to an additional ALE experiment based on the results of shake flask fermentations. MT01 showed enhanced tolerance to acetic acid compared to the parent since it was able to ferment xylose in the highest concentration of acetic acid tested in shake flasks containing YPXAC. However, this performance was not observed in SBHH because MT01 did not grow in hydrolysate. It was not surprising because the hydrolysate had high amounts of acetic acid, as well as a mixture of other inhibitors, which likely had a synergistic effect. After ALE, the evolved strain ME3.5.5 could ferment xylose to ethanol in SBHH. Interestingly, no trade-offs were observed for the strain ME3.5.5. Trade-offs are common in ALE experiments, and strains that do not show this feature are interesting for biotechnological purposes (Dragosits and Mattanovich, 2013; Van den Bergh et al., 2018).

In ASGH, the evolved strain exhibited similar growth and metabolite production to the parental strain, likely due to the lower concentration of inhibitors and the higher availability of sugars compared to SBHH. A hypothesis is that the evolved strain has adaptations that enhance its tolerance to inhibitors present in SBHH, allowing it to grow where the parental strain cannot. However, in the more favorable ASGH, with fewer inhibitors and more abundant glucose and xylose, both grow at similar rates, as the evolved strain’s enhanced inhibitor tolerance may no longer provide a significant advantage. Thus, the availability of sugars and reduced stress conditions likely neutralize any metabolic differences between the strains.

A major goal of this study was to map the genetic basis of improved growth and fermentation in hydrolysate conditions. Mutagenesis of UFMG-CM-Y469 led to strain MT01, which showed improved acetic acid tolerance. Therefore, the improved phenotype of evolved strain ME3.5.5 in YPXAC is presumably due to additive contributions of mutations acquired in the original mutagenesis of UFMG-CM-Y469 and those acquired during the ALE of MT01. Five mutations were identified in ME3.5.5, four of which are predicted to disrupt in protein-coding sequences. Because we did not sequence MT01, we cannot hypothesize which of these mutations underlie the initial improved acetic acid tolerance of MT01 and which mutations drove the adaptive increase in acetic acid tolerance in ME3.5.5.

The four genes predicted to contain disruptive heterozygous mutations in ME3.5.5 are thus candidates for genetic modification to increase growth in YPXAC and SBHH. These genes include one hypothetical protein-encoding gene and three genes conserved in *S. cerevisiae*. Homologs of the *S. cerevisiae* genes *ERT1* (36% amino acid identity), *NPL4* (49% amino acid identity), and *CYR1* (43% amino acid identity) acquired a nonsynonymous, nonsense, and frameshift mutations, respectively. Of these, *CYR1* has a previously established connection to xylose fermentation (Wu et al., 2020). *CYR1* encodes adenylate cyclase, which converts AMP to cAMP in response to glucose stimulation of Gpr1p. Elevated levels of cAMP then activate the PKA pathway, upregulating glycolysis and downregulating gluconeogenesis. Elevated cAMP and associated elevation in PKA signaling have been implicated in increased ability to ferment xylose in *S. cerevisiae* (Sato et al., 2016; Wu et al. 2020). However, the frameshift we observed is predicted to truncate the protein and eliminate the C-terminal catalytic domain. Mutations to this domain have been previously shown to inactivate Cyr1p and correspondingly decrease cAMP levels (Vanhalewyn et al., 1999). Nonetheless, the truncation we observed was heterozygous, so it may only decrease adenylate cyclase activity by half. However, we also hypothesize that it could also be dominant negative, meaning the truncated mutant protein may interfere with the function of the wild-type allele, leading to a more severe reduction in activity.

The parental strain used (UFMG-CM-Y469) is a Brazilian isolate and not derived from the reference genome isolate from the United States of America (NRRL Y-27907). We considered that additional mutations may be obscured in regions unique to UFMG-CM-Y469, but we did not find any additional variants when using a mapping-free approach (Standage et al., 2019).

## Conclusion

The combination of random mutagenesis and adaptive laboratory evolution (ALE) were effective tools for selecting mutant *Spathaspora passalidarum* strains with enhanced fitness under stressful conditions, such as acetic acid exposure and sugarcane bagasse hemicellulosic hydrolysate (SBHH). The mutant generated by ultraviolet-induced mutagenesis and subsequent evolution experiments showed improved growth and ethanol production over the parental strain in these challenging environments. The identification of a heterozygous mutation in the adenylate cyclase-encoding gene, *CYR1*, in the evolved strain suggests a potential role in enhancing fitness, as evidenced by its known role in glucose response and fermentation in *S. cerevisiae*. However, the lack of unique mutations in MEH30.1 and the similar fitness observed in evolved strains in SBHH likely indicate that the additional ALE did not select for new mutants. Further investigation,particularly into cAMP signaling in the mutants, will be necessary to fully understand the mechanisms driving adaptation in these strains.

## Data availability

The raw data (NCBI’s SRA) will be released upon publication.

## Figure’s Legend

Figure S1: Shake flask fermentations of YPX (5% xylose) in 72 h by *Sp. passalidarum* ME3.5.5. Error bars indicate the standard deviation from triplicate shake flasks.

Figure S2: Shake flask fermentations of sugarcane bagasse hemicellulosic hydrolysate (SBHH) in 96 h by *Sp. passalidarum* MEH30.1. Error bars indicate the standard deviation from triplicate shake flasks. AC, acetic acid.

