## Supplementary material for "An evolved strain of *Spathaspora passalidarum* produces ethanol from sugarcane bagasse and switchgrass lignocellulosic hydrolysates": FigureS1_and_FigureS2

[1]Departamento de Microbiologia, ICB, C.P. 486, Universidade Federal de Minas Gerais, Belo Horizonte, Brazil.

[2]Laboratory of Genetics, J. F. Crow Institute for the Study of Evolution, Wisconsin Energy Institute, Center for Genomic Science Innovation, University of Wisconsin-Madison, Madison, WI 53726, USA.

[3]DOE Great Lakes Bioenergy Research Center, University of Wisconsin-Madison, Madison, WI 53726, USA.

[4]Department of Biotechnology, Engineering School of Lorena, University of São Paulo, Estrada Municipal do Campinho s/n, Lorena, SP, 12602-810, Brazil.

*Deceased

#Correspondence:

Chris Todd Hittinger

Carlos A. Rosa

### Abstract

Lignocellulosic hydrolysates, derived from plant biomass, contain various inhibitors that can hinder microbial growth. This study aimed to enable the growth and ethanol production by the xylose-fermenting yeast *Spathaspora passalidarum* in the presence of lignocellulosic hydrolysate inhibitors, particularly acetic acid. Ultraviolet (UV)-induced mutagenesis and adaptive laboratory evolution (ALE) were used to select for mutants with higher tolerance to these inhibitors. The initial mutant strain, MT01, was selected for increased growth in medium containing xylose and acetic acid. This strain underwent further evolution, resulting in the strain ME3.5.5, which showed significant improvements in both growth and ethanol production compared to the parental strain when tested in sugarcane bagasse hemicellulosic hydrolysate (SBHH). Genomic analysis identified non-synonymous and frameshift mutations in four genes, including *CYR1* (encoding adenylate cyclase). These findings suggest that genetically optimized *S. passalidarum* strains could play a crucial role in advancing industrial bioethanol production from lignocellulosic biomass by overcoming the inhibitory effects of compounds found in lignocellulosic hydrolysates.

Keywords: mutagenesis, Adaptive Laboratory Evolution (ALE), *Spathaspora passalidarum*, acetic acid, hemicellulosic hydrolysates.

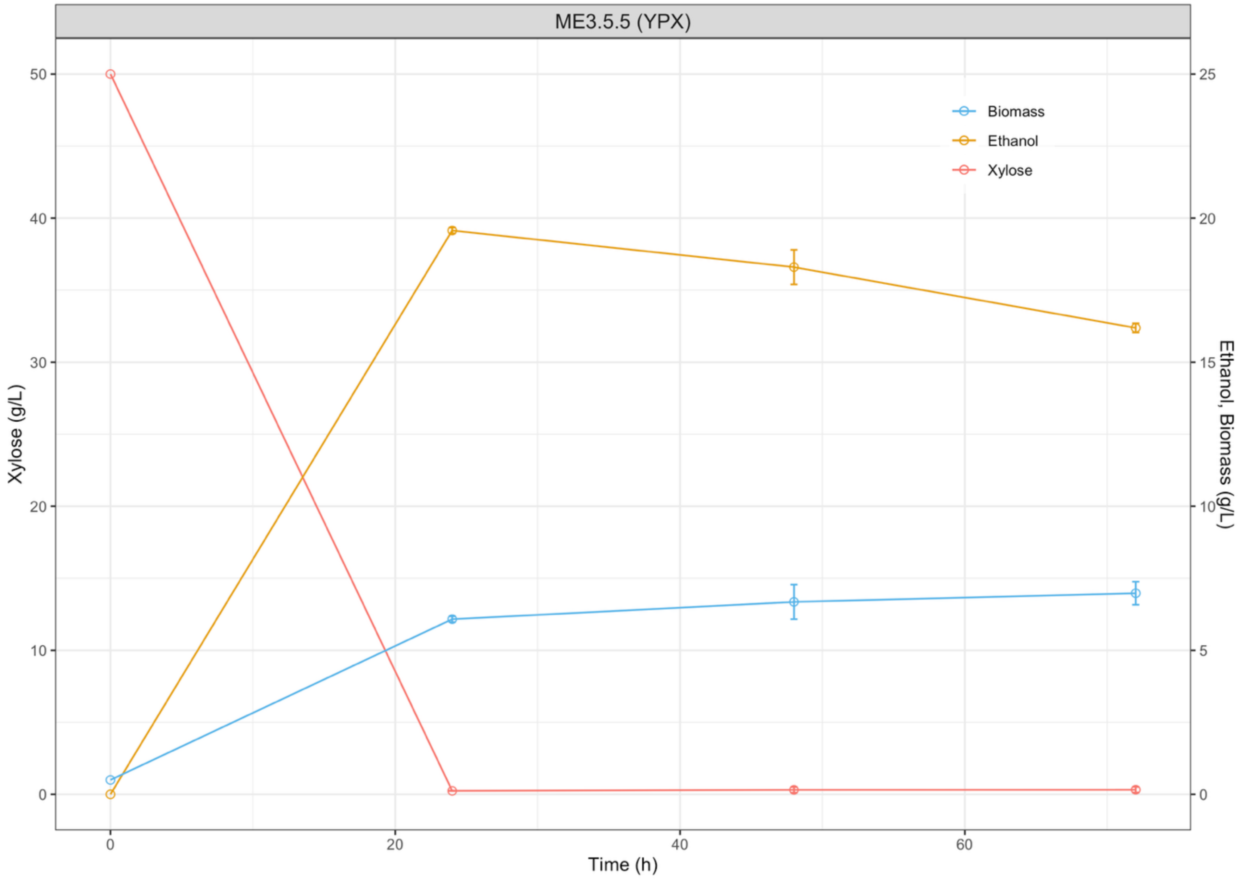


Figure S1: Shake flask fermentations of YPX (5% xylose) in 72 h by *Sp. passalidarum* ME3.5.5. Error bars indicate the standard deviation from triplicate shake flasks.

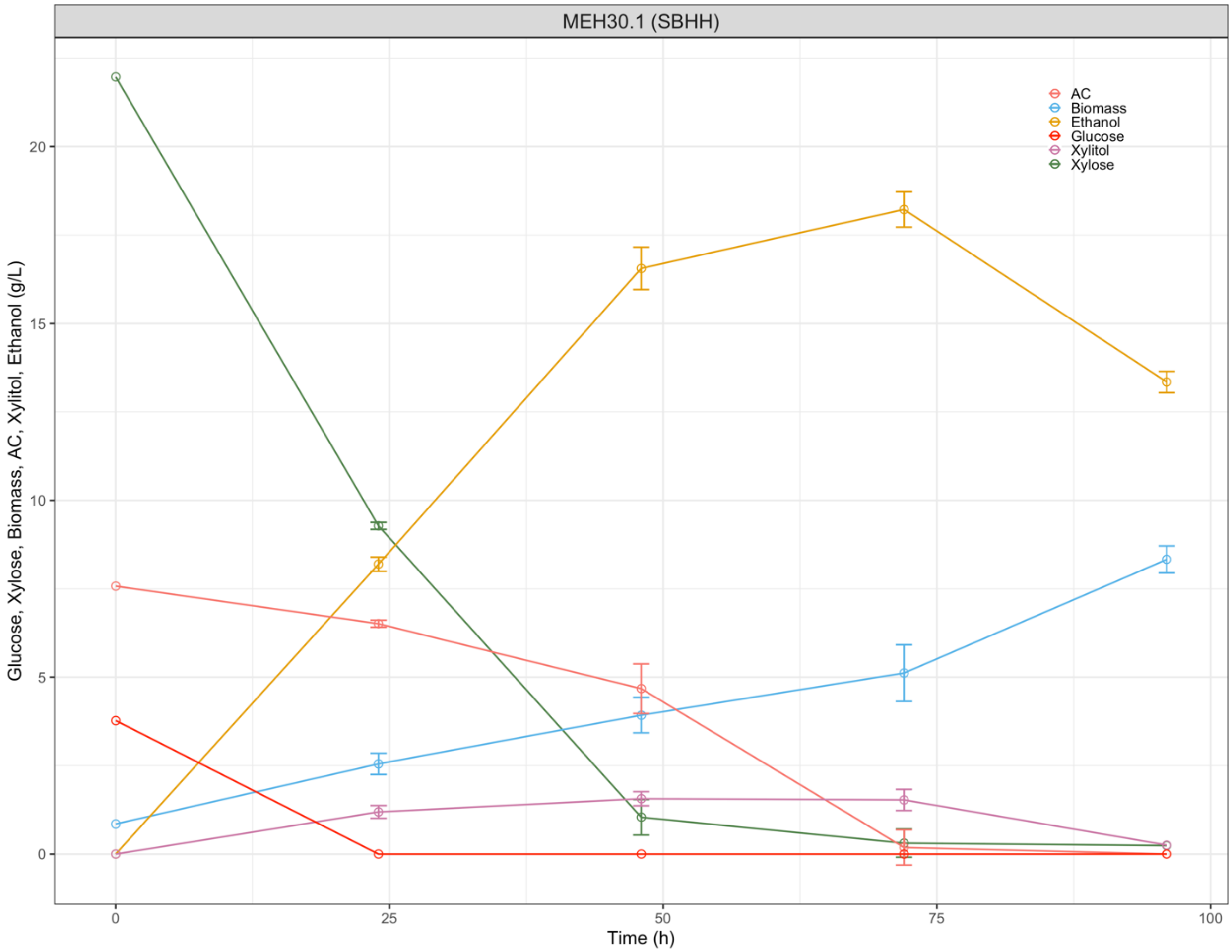


Figure S2: Shake flask fermentations of sugarcane bagasse hemicellulosic hydrolysate (SBHH) in 96 h by *Sp. passalidarum* MEH30.1. Error bars indicate the standard deviation from triplicate shake flasks. AC, acetic acid.
